# A novel regulatory gene promotes novel cell fate by suppressing ancestral fate in the sea anemone *Nematostella vectensis*

**DOI:** 10.1101/2021.08.29.458124

**Authors:** Leslie S Babonis, Camille Enjolras, Joseph F Ryan, Mark Q Martindale

## Abstract

Cnidocytes (“stinging cells”) are an unequivocally novel cell type used by cnidarians (corals, jellyfish, and their kin) to immobilize prey. Although they are known to share a common evolutionary origin with neurons, the developmental program that promoted the emergence of cnidocyte fate is not known. Using functional genomics in the sea anemone, *Nematostella vectensis*, we show that cnidocytes evolved by suppression of neural fate in a subset of neurons expressing RFamide. We further show that a single regulatory gene, a C_2_H_2_-type zinc finger transcription factor (ZNF845), coordinates both the gain of novel (cnidocyte-specific) traits and the inhibition of ancestral (neural) traits during cnidocyte development and that this gene arose by domain shuffling in the stem cnidarian. Thus, we uncover a mechanism by which a truly novel regulatory gene (ZNF845) promoted the origin of a truly novel cell type (cnidocyte) through duplication of an ancestral cell lineage (neuron) and inhibition of its ancestral identity (RFamide).

**Significance:** In this study, we demonstrate how new cell types can arise in animals through duplication of an ancestral (old) cell type followed by functional divergence of the new daughter cell. Specifically, we show that stinging cells in cnidarians (jellyfish and corals) evolved by duplication of an ancestral neuron followed by inhibition of the RFamide neuropeptide it once secreted. This is the first evidence that stinging cells evolved from a specific subtype of neurons and suggests some neurons may be easier to co-opt for novel functions than others.

## Introduction

Understanding the mechanisms driving cell type diversification persists as one of the key challenges in evolutionary biology (*1*). The gain of new adaptive cell functions requires either the advent of novel genes (*2*, *3*). the modification of existing gene regulatory networks (*4*), or some combination of these two processes (*5*). Alone, this additive model, focused simply on the emergence of novel gene interactions, is insufficient to explain expansion of cell identity as new cell types would arise in place of ancestral cell types. In a process analogous to gene duplication and divergence (*6*), new instances of cell division during embryogenesis could lead to duplication of a cell lineage, providing the opportunity for one lineage to retain an ancestral function and the other to acquire new functions. Cell type diversification, therefore, requires both novel gene interactions and a novel cell lineage in which to express these traits.

Cnidarians are an unparalleled model for studying the evolution of novelty because the defining synapomorphy of this group (the cnidocyte or “stinging cell”) is an unequivocally novel cell type. During embryogenesis, cnidocytes differentiate from a progenitor cell that also gives rise to neurons, reflecting a common evolutionary origin for these two cell types (*7*, *8*). Two key features were necessary for the transition away from neural fate in early cnidocytes: the development of the explosive secretory organelle (the cnido<u>cyst</u>) from which cnidocytes derive their “sting,” and the suppression of neural phenotype (axons, synaptic signaling molecules, etc). Studies tracking the synthesis of the cnidocyst-specific protein minicollagen have revealed key steps leading to the origin of this novel organelle (*9*, *10*); the mechanisms driving the suppression of neural phenotype in cnidocytes remain unknown. Here, we show that a single novel transcription factor, ZNF845, both promotes cnidocyte fate and suppresses neural fate during development of the sea anemone *Nematostella vectensis*. We further show that the six-domain topology of ZNF845 arose through domain shuffling in the last common ancestor of cnidarians, making this a clear example of a cnidarian-specific gene driving development of a cnidarian-specific trait.

## Results

### ZNF845 promotes cnidocyte fate

Beginning at the blastula stage, ZNF845 is expressed throughout the embryo in cells that are actively undergoing DNA synthesis, as labeled by EdU (**Fig. 1A**). ZNF845 is co-expressed with SoxB2 in a subset of embryonic neural progenitor cells at the blastula stage (**Fig. 1B**) and is also co-expressed in a subset of PaxA-expressing developing cnidocytes at the gastrula stage (**Fig. 1C**). ZNF845 continues to be expressed through metamorphosis in a pattern reminiscent of cnidocyte development (Supplementary Material Fig. S1). To understand the role of ZNF845 in neural/cnidocyte differentiation, we knocked down SoxB2 and PaxA using previously published morpholinos (MOs) (*8*, *11*). ZNF845 expression was downregulated in embryos injected with the SoxB2 MO but unaffected in embryos injected with PaxA MO (**Fig. 1D**). Paired with the co-expression results, these data suggest that ZNF845 is part of the cnidogenesis pathway acting upstream of PaxA in *N. vectensis*.

**Fig. 1.**
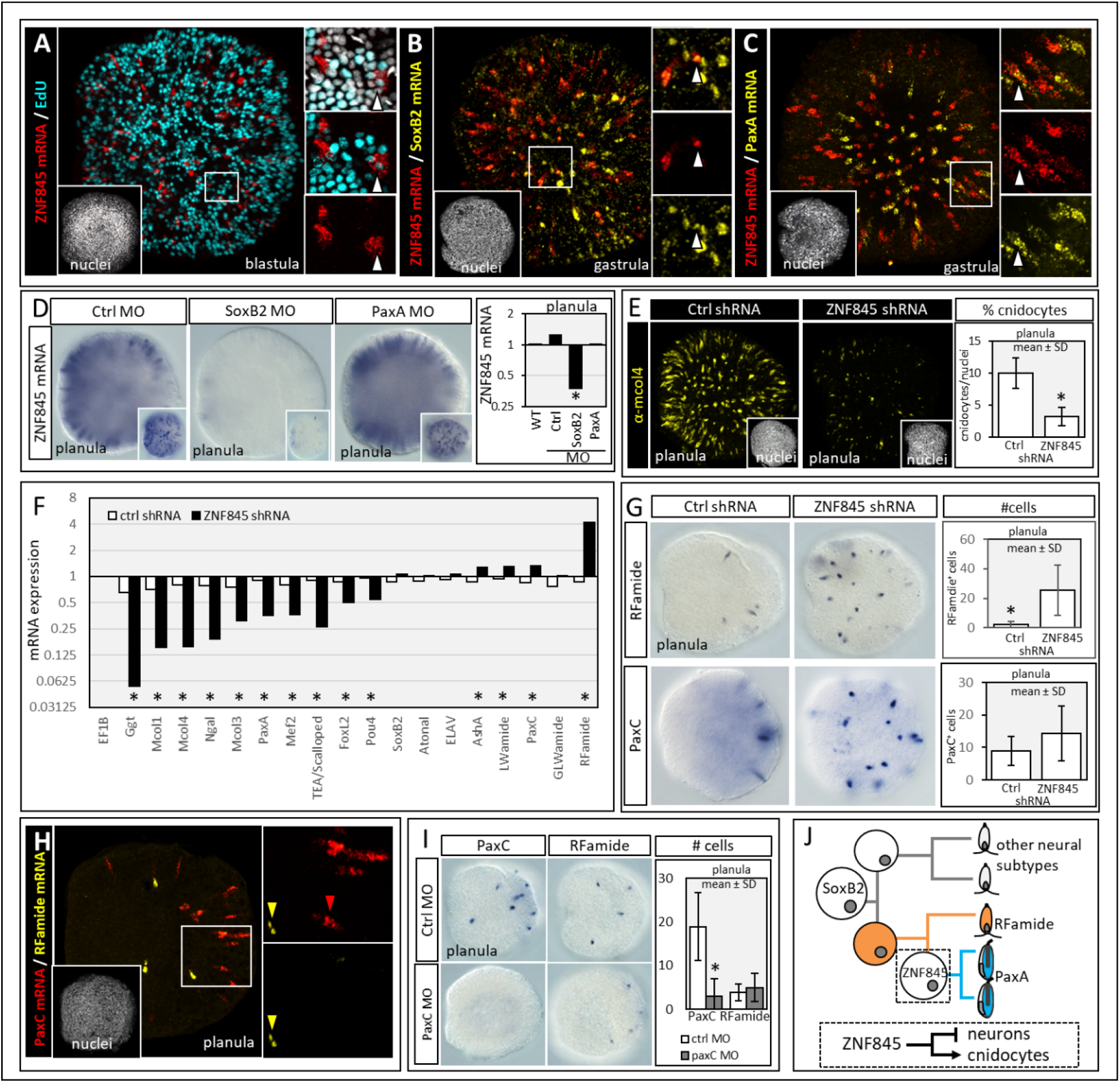
ZNF845 specifies cnidocyte identity in *N. vectensis*. (A-C) ZNF845 is partially co-expressed with: (A) EdU (proliferating cells), (B) SoxB2 mRNA (neural progenitor cells), and (C) paxA mRNA (cnidocytes). Insets: white arrowheads show co-expression; nuclei are white (DAPI). (D) ZNF845 expression after knockdown of SoxB2 and PaxA (by morpholino, MO) assayed by *in situ* hybridization and qPCR; significance (*) is indicated for SoxB2MO or PaxA MO vs Ctrl MO. (E) Cnidocyte differentiation (α-Mcol4 antibody) after ZNF845 knockdown (shRNA). (F) qPCR of target gene expression after ZNF845 knockdown; fold-change relative to housekeeping gene EF1B. (G) Response of RFamide- and PaxC-expressing cells to ZNF845 knockdown. (H) RFamide and PaxC are not co-expressed. (I) Expression of RFamide and PaxC after knockdown of PaxC (MO). (J) Model for ZNF845-mediated specification of cnidocyte identity from and RFamide-expressing cell lineage. Significance (*) for all tests indicated as p<1E-02. See Table S2 for supporting information.

To further explore the role of ZNF845, we knocked down ZNF845 using small-hairpin RNAs (shRNAs) (*12*) and assayed the effects on cnidocyte development. Knockdown of ZNF845 was effective through the late planula stage (Supplementary Material Fig. S1) and resulted in nearly complete loss of cnidocytes throughout the ectoderm of the planula (**Fig. 1E**). Using an antibody directed against the cnidocyte-specific protein minicollagen4 (α-Mcol4) (*13*) we demonstrate a significant loss of cnidocytes: from 10% of total cells in wildtype (WT) and control (ctrl) shRNA-injected embryos to less than 3% when ZNF845 was knocked down. We recovered identical results in embryos injected with a splice-blocking ZNF845 MO, relative to those injected with a standard control MO (Supplementary Material Fig. S2).

To determine if ZNF845 acts upstream of PaxA and other markers specific to the cnidogenesis pathway, we examined the effect of ZNF845 knockdown on known markers of neural and cnidocyte differentiation in *N. vectensis* using quantitative PCR (**Fig. 1F, Table S2**). Knockdown of ZNF845 resulted in significant downregulation of PaxA, and all of its known targets (GGT, Ngal, Mcol1, Mcol3, Mcol4, and Mef2IV) (*11*) as well as three cnidocyte-expressed transcription factors identified from single-cell RNA-Seq analysis (TEA/Scalloped, FoxL2, Pou4) (*14*). Conversely, ZNF845 knockdown did not affect the expression of SoxB2 or the neuron-specific regulatory genes Atonal and ELAV (*15*). While ZNF845 knockdown caused a statistically significant increase in the expression of neural markers AshA, GLWamide, LWamide, and PaxC assayed by qPCR, the response of these genes was minor relative to the large, significant upregulation of RFamide. To spatially characterize these results, we performed *in situ* hybridization for RFamide and PaxC in embryos injected with control- or ZNF845 shRNAs and counted the number of cells expressing these neural markers at the gastrula stage (**Fig. 1G**). Knockdown of ZNF845 significantly increased the number of RFamide-expressing cells from 2.23 (± 1.95; mean ± SD) in control embryos to 23.47 (± 17.85; mean ± SD) in ZNF845 knockdowns. Cell counts for PaxC expression revealed a small, non-significant increase in the number of PaxC-expressing cells from 8.9 (± 4.4; mean ± SD) in controls to 14.3 (± 18.5; mean ± SD) in ZNF845 knockdowns. Because PaxC expression during embryogenesis is reminiscent of RFamide expression, we hypothesized that PaxC might be an upstream regulator of RFamide neuron differentiation. On the contrary, we found that PaxC and RFamide are not co-expressed (**Fig. 1H**) and that knockdown of PaxC using a previously published MO (*11*) did not affect the number or distribution of RFamide-expressing cells (**Fig. 1I**).

These results suggest that there is a closer evolutionary relationship between cnidocytes and RFamide-expressing neurons than between cnidocytes and other neural subtypes (**Fig. 1J**). This is congruent with a recent study showing that the selector gene Pou4 regulates terminal cell identity in cnidocytes and RFamide-expressing neurons in *N. vectensis* but not in other cell types (*16*). Second, we demonstrate that a single transcription factor (ZNF845) upregulates both the genes necessary to promote cnidocyte identity and the genes necessary to inhibit RFamide neuron identity. To understand the mechanism by which ZNF845 suppresses RFamide expression during cnidocyte differentiation, we searched for inhibitory transcription factors that were co-expressed with cnidocyte specific genes.

### ZNF845 inhibits neural fate through NR12

Nuclear receptors in the COUP-TF family (NR2F) are known to play an inhibitory role in neural cell fate decisions in both cnidarians (*17*) and bilaterians (*18*). In *N. vectensis*, there are five NR2F paralogs: NR10-14, most of which appear to have originated through lineage-specific duplication in cnidarians (*19*). We examined the expression of all five NR2F genes (Supplementary Material Fig. S3) and found that three of them (NR11,12,13) were expressed in the ectoderm during early embryogenesis and were also downregulated in embryos injected with ZNF845 shRNA (**Fig. 2A**). All three were also co-expressed with Mcol4 in differentiating cnidocytes (**Fig. 2B-D**) and yet we found no evidence of that any of the NR2F paralogs were co-expressed together in the same cell (**Fig. 2E,F**) suggesting these NR2F paralogs are expressed uniquely in different subpopulations of cnidocytes.

**Fig. 2.**
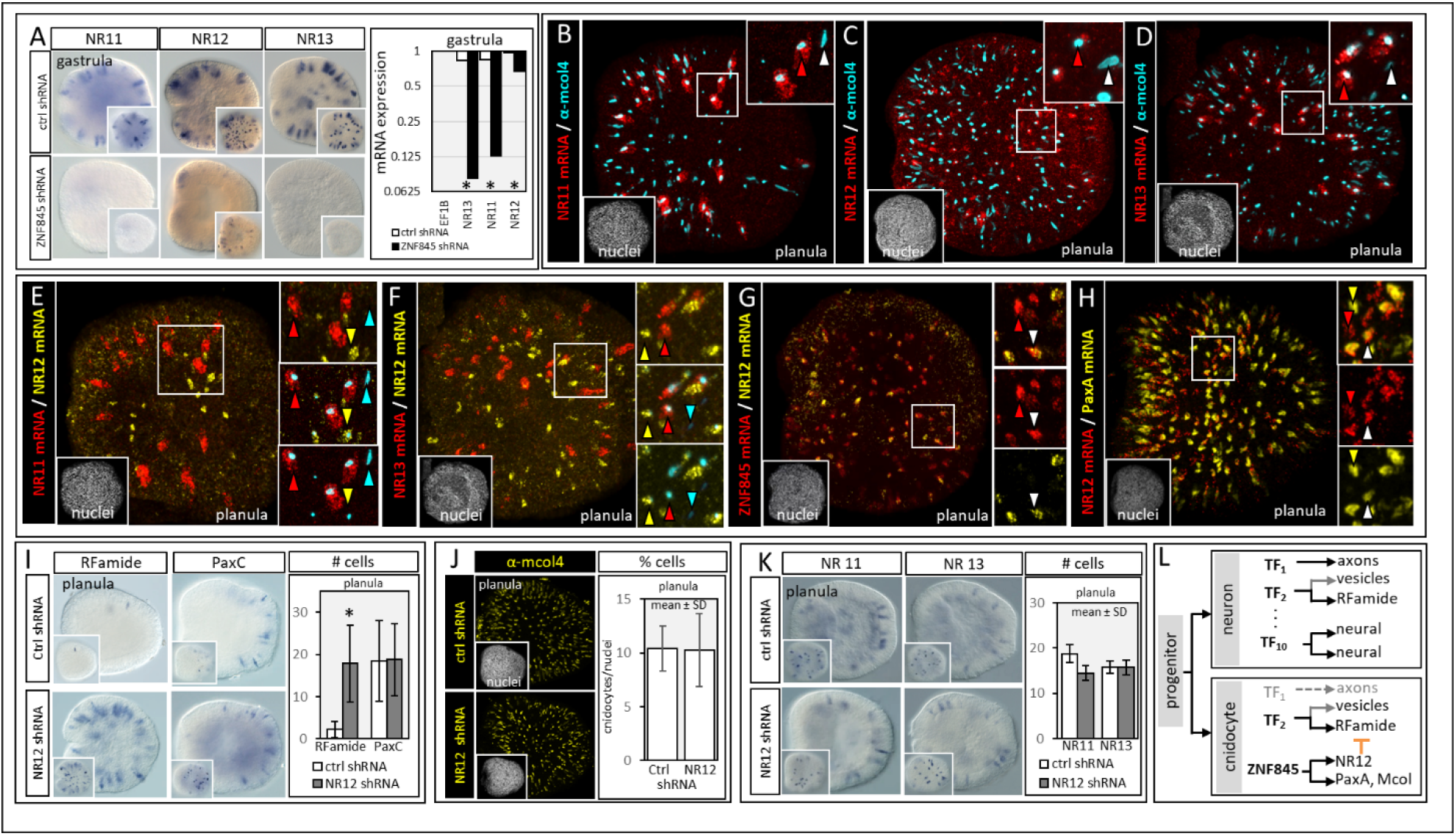
RFamide identity is inhibited by NR12 in developing cnidocytes. (A) NR11, NR12, and NR13 are downregulated in ZNF845 knockdowns (shRNA) as assayed by *in situ* hybridization and qPCR. Insets: surface detail. (B-D) NR11, NR12, and NR13 are co-expressed with Mcol4 (cnidocytes), but not with each other (E,F). Images B and E show the same embryo, as do D and F. NR12 is partially co-expressed with (G) ZNF845 and (H) PaxA. (I) Response of RFamide- and PaxC-expressing cells after NR12 knockdown. No effect of NR12 knockdown on (J) cnidocyte differentiation (α-Mcol4) or (K) expression of NR11 or NR13. (L) Modular regulation of neural traits (e.g., axons, vesicles + neuropeptides) explains how some traits can be lost (e.g., axons) and others retained but inhibited (RFamide) in developing cnidocytes. Significance (*) for all tests indicated as p<1E-02. See Table S2 for supporting information.

The cnidocyte-specific expression of NR12 that we observed was also supported by single cell RNA-Seq in *N. vectensis*(*14*). We examined NR12 further and found that it is co-expressed in a subset of ZNF845-expressing cells at the gastrula stage (**Fig. 2G**) and in a subset of PaxA-expressing cells at the early planula stage (**Fig. 2H**). We then examined the effects of NR12 on cnidocyte development using shRNAs (Supplementary Material Fig. S4, Table S1) and found that knockdown of NR12 resulted in a four-fold increase in the number of RFamide-expressing cells but had no effect on expression of PaxC (**Fig. 2I**) or on the specification of cnidocytes (**Fig. 2J**). Knockdown of NR12 also had no effect on the number or distribution of NR11- or NR13-expressing cells (**Fig. 2K**).

Modularity in the regulation of neural phenotype would have enabled cnidocytes to retain beneficial aspects of the ancestral phenotype while silencing others through selective inhibition, as has been shown for neural subtype specification in C. elegans (*18*). The upregulation of RFamide following NR12 knockdown suggests that RFamide expression may be coupled to the expression of another trait that had adaptive value during the evolution of cnidocytes (e.g., secretory vesicles) (**Fig. 2L**). The independent expression of NR11 and NR13 in non-overlapping populations of developing cnidocytes suggests that multiple cnidocyte subtypes may be specified through NR2F-mediated inhibition of neural traits in other (non-RFamide) neural subtypes.

### ZNF845 is a novel transcription factor

Transcription factors with Cys2-His2 zinc finger (ZF-C_2_H_2_) domains (PF00096) represent one of the largest families of transcription factors across animals (*20*). Numerous evolutionary processes have contributed to diversification in this gene family, including duplication/divergence, gain/loss of ZF-C_2_H_2_ domains, and gain/loss of accessory domains (*21*). Among the oldest members of this clade are ZNF proteins with multiple tandem C_2_H_2_ domains but no other conserved functional domains (*22*). ZNF845 has six tandem ZF-C_2_H_2_ domains; to understand how these six domains came together in a single protein, we generated a maximum likelihood phylogeny of all ZF-C_2_H_2_ domains extracted from predicted proteomes for three bilaterian taxa (*Homo sapiens*, *Drosophila melanogaster*, and *Caenorhabditis elegans*) and four cnidarians (two anthozoans: *N. vectensis* and *Acropora digitifera*, and two medusozoans: *Hydra magnipapillata* and *Nemopilema nomurai*). The full phylogeny contains over 11,000 branch tips; the FASTA alignment is provided in Supplementary Material Data S1.

*Nematostella vectensis* has 218 predicted proteins with one or more ZF-C_2_H_2_ domains, sixteen of which, including ZNF845, encode six tandem ZF-C_2_H_2_ domains. We examined the evolutionary relationships of the ZF-C_2_H_2_ domains from ZNF845 (JGI protein ID: 81344) and compared the results with an analysis of another six-domain ZNF protein, Growth Factor Independence 1B (Gfi1B; JGI PID: 112378). Gfi1B is a potent regulator of cell differentiation in vertebrates (*23*), but the function of this protein has not been characterized in *N. vectensis*. Each of the six domains from the *N. vectensis* ortholog of ZNF845 groups with ZF-C_2_H_2_ domains from the orthologous ZNF845 protein in other cnidarians: Hmag_XP_002158383.2 (originally named ZNF845 by Hemmrich et al. (*24*)), Nnom_10870, and Nnom_3755 (**Fig. 3A** and Supplementary Material Fig. S5). These relationships suggest the anthozoan and medusozoan ZNF845 proteins descended from a common ancestor that already had six ZF-C_2_H_2_ domains in the stem cnidarian. Additionally, two of these domains (domains 1 and 5) appear to have arisen by tandem duplication as these domains form a clade that lacks other ZF-C_2_H_2_ domains. Each of the six ZF-C_2_H_2_ domains in ZNF845 shares some level of homology with a bilaterian ZF-C_2_H_2_ domain, but none of these bilaterian domains are found in the same protein.

**Fig. 3.**
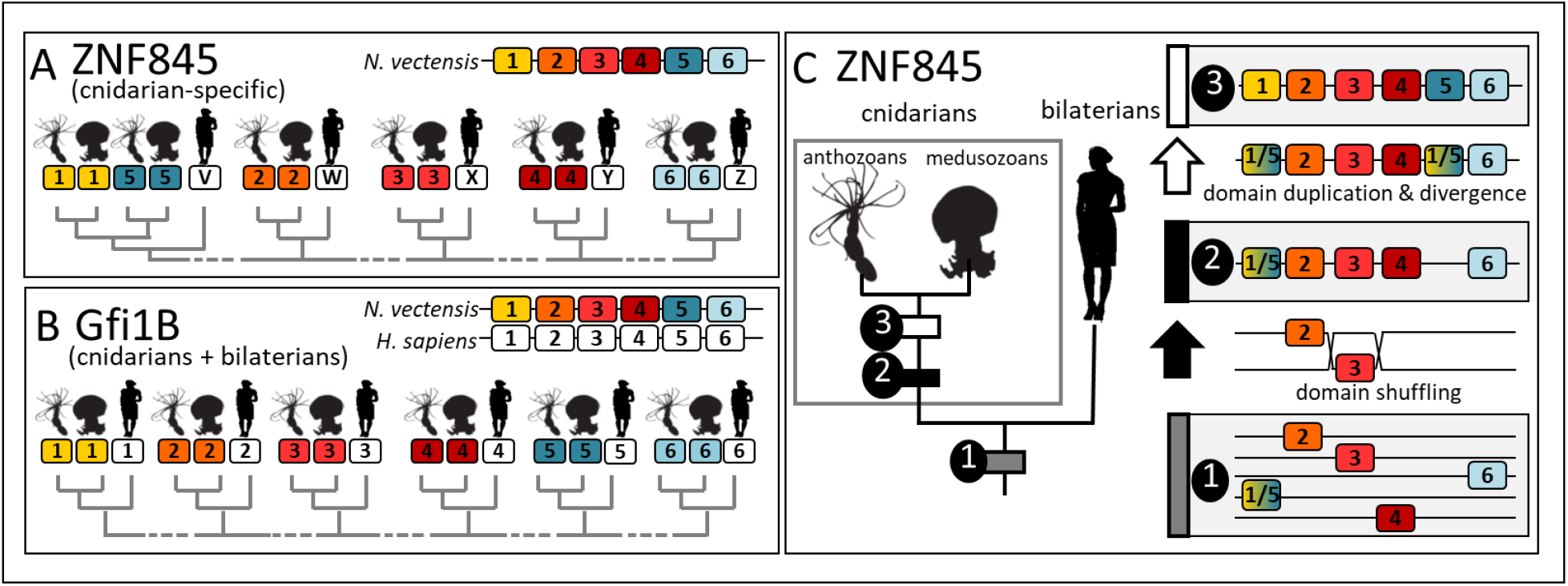
Evolutionary history of two 6-domain ZNF proteins. Relationships among ZF-C_2_H_2_ domains from (A) ZNF845 (*N. vectensis* JGI PID 81344) and (B) Gfi1B (*N. vectensis* JGI PID: 112378). (C) Proposed model for the emergence of ZNF845: 1. In the common ancestor of cnidarians and bilaterians ZF-C_2_H_2_ domains were in distinct proteins, 2. In the stem cnidarian, domain shuffling brought domains 1/5, 2, 3, 4 and 6 together in a single protein, 3. Domain 1/5 then duplicated and diverged to become two distinct domains (1 and 5) before the diversification of extant cnidarians. Broken grey lines in A, B represent the locations of the other ~11,000 branch tips of the complete tree. See Supplementary Figs S5,S6 for supporting information.

Examination of the ZF-C_2_H_2_ domains from Gfi1B suggests that, unlike ZNF845, this six-domain transcription factor emerged in its current form before cnidarians and bilaterians diverged from their common ancestor (**Fig. 3B**, Supplementary Material Fig. S6). Each of the ZF-C_2_H_2_ domains from the *N. vectensis* ortholog of Gfi1B grouped with the syntenic ZF-C_2_H_2_ domain from the Gfi1B ortholog in each of the bilaterian taxa examined (Homo_NP_001120687.1, Homo_XP_006717360.1, Dmel_Q9VM77, Cele_5376). Together, these observations suggest that ZNF845 arose as a novel 6-domain protein in the stem cnidarian and that both domain shuffling and domain duplication/divergence were important for the emergence of this protein (**Fig. 3C**).

### Modeling cell type expansion

Gene duplication and divergence is an important source for novel gene function (*6*, *25*). Extending this concept, we propose a model for the emergence of novel cell types through cell lineage duplication and divergence (**Fig. 4**). New cells are generated from progenitors during growth and tissue repair (**Fig. 4A**). Co-option of a progenitor cell to produce a novel daughter would not increase the number of differentiated cell types if the novel cell fate simply replaced its ancestor (**Fig. 4B**). Duplication of the progenitor cell first would allow for maintenance of ancestral cell identity and the origin of novel identity, increasing cell type diversity (**Fig. 4C**).

**Fig. 4.**
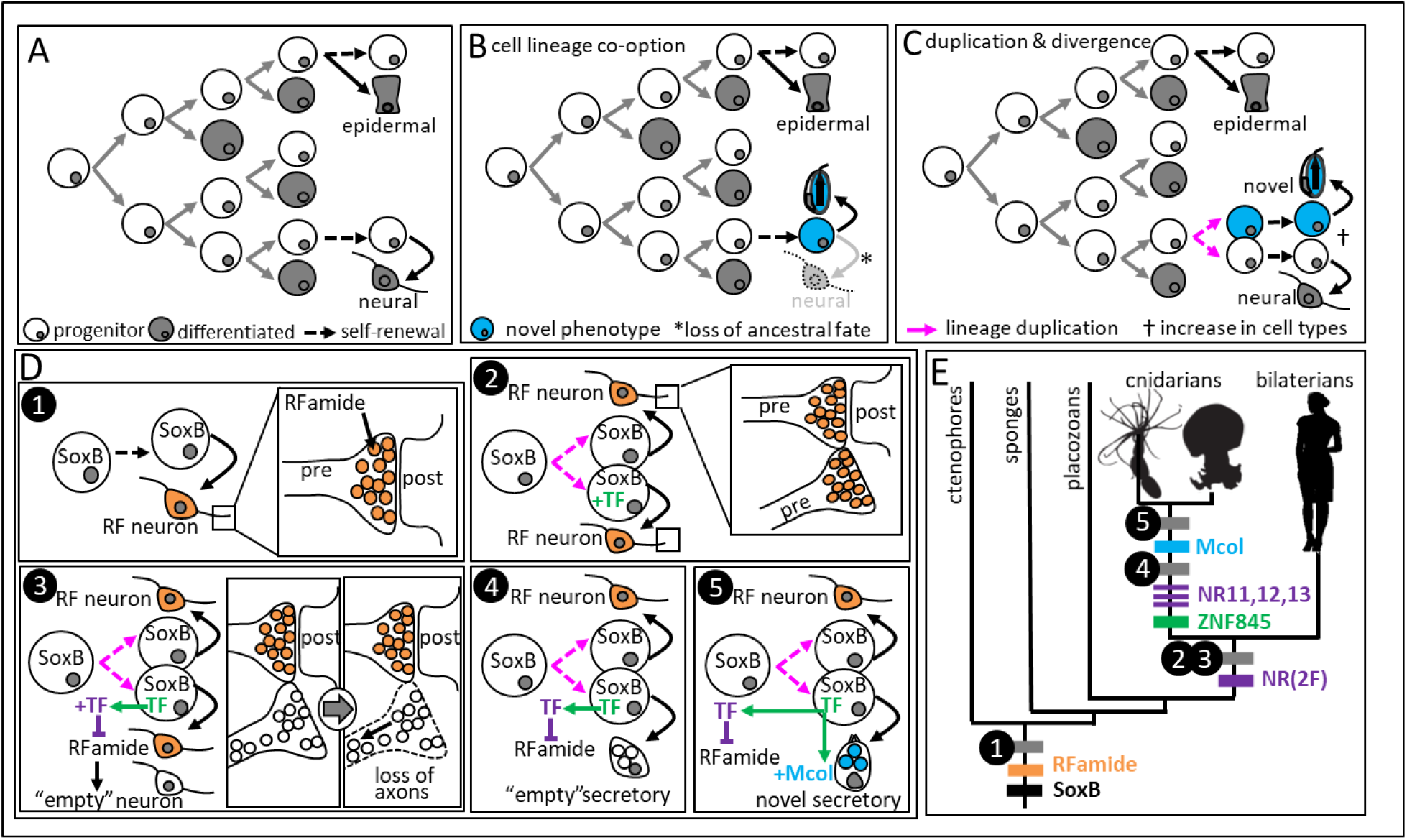
Origin of novelty through lineage duplication and suppression of ancestral fate. (A) Embryogenesis as a model for cell type evolution. (B) Co-option of a progenitor cell for novel fate does not increase cell type diversity. (C) Duplication of the progenitor cell prior to co-option allows for the expansion of differentiated cell types. (D) A stepwise model for the origin of cnidocytes by duplication and divergence of an RFamide neural lineage. (E) A proposal for the evolutionary timing of these events; similar duplication of a neural progenitor cell and inhibition of the neural payload (steps 2,3) may also have facilitated the expansion of neural subtypes in early bilaterians.

These basic principles can be extended to explain the emergence of cnidocytes from a neural precursor (**Fig. 4D,E**). At least one SoxB gene was present in the ancestor of all animals (*26*). Analyses of animal neural diversity have suggested RFamide-like neurons arose early in animal evolution as well (*27*, *28*). To generate a novel cell lineage, the progenitor cell would first have to duplicate to generate a new progenitor lineage. Absent additional changes, duplication of a neural progenitor cell would have doubled the number of RFamide-expressing daughters, potentially leading to aberrant synapse formation. If progenitor cell duplication were coupled to the origin of a transcription factor that inhibited RFamide expression, the original number of RFamide-expressing daughters would be maintained and RFamide neurons would now be sister to a second lineage of “neurons” lacking a synaptic payload. Key to the emergence of cnidocytes, therefore, was the origin of two transcription factors: one (ZNF845) that could maintain progenitor cell fate in a newly duplicated cell lineage and another (NR12) that could manipulate the identity of the secreted payload in the duplicated daughter cell. Without the selection pressure to maintain synaptic signaling in this new daughter cell, these cells may have experienced relaxed selection for the maintenance of axons, allowing the secretory vesicles to relocate to the cell body. Subsequent mutations resulting in the emergence of a novel payload (e.g., minicollagen) would have further promoted daughter cell divergence following duplication in this cell lineage.

## Discussion

The stepwise model presented here illustrates three important paradigms for the evolution of novel cell types. First, although the role of novel genes in driving evolutionary innovation has been debated (*29*–*31*), cnidocytes have always provided a clear example of an adaptive role for novel effector genes (e.g., minicollagen) in driving the evolution of a novel cell phenotype (*9*). In the present study, we extend this adaptive role for novel genes up the cnidocyte gene regulatory network by showing that the emergence of a new transcription factor assembled through domain shuffling in the stem cnidarian (ZNF845) was essential for the origin of cnidocytes.

Second, we demonstrate support for the hypothesis that modularity is an important driver of phenotypic evolution (*32*, *33*). During the early divergence of cnidocytes, modularity in neural cell phenotype allowed for selective inhibition of certain traits (e.g., vesicular payload, axons) and retention of others (e.g., secretory vesicle). In this scenario, a single mutation that allowed for inhibition of RFamide expression could rapidly change the selection pressures affecting the recently duplicated sister cells, promoting retention of ancestral phenotype in one and allowing additional mutations to arise in the other.

Finally, our results broadly support the hypothesis that the origin of a secretory cell lineage was advantageous for the emergence of diverse novel cell functions (*34*, *35*). The ability to segregate gene products into a compartment within the cell and to target those products for delivery to the extracellular space allows for the retention of novel gene products that may otherwise be deleterious if retained intracellularly (e.g., collagen fibers).

Applied broadly, this scenario of secretory cell duplication coupled to payload inhibition combined with the early diversification of novel neuropeptides (*36*–*38*), could also explain the rapid expansion of neural function in early bilaterians. Studies of neural fate in bilaterian model systems have demonstrated that discrimination of specific subtypes within a neural lineage relies on inhibition of the effector genes that define other subtypes in the lineage (*39*), and that this inhibition of sister cell fate is mediated through the actions of conserved transcription factors including orthologs of NR2F/COUP-TF (*18*). The data presented here indicate that this role for NR2F-mediated inhibition of cell fate may extend back to the common ancestor of cnidarians and bilaterians, suggesting the divergence-through-inhibition regulatory logic was already driving the expansion of cell fate nearly 700 million years ago (*40*).

## Materials and Methods

### Gene knockdowns

To assess the influence of ZNF845 and NR12 on cnidocyte fate, we performed mRNA knockdown by microinjection of small-hairpin RNAS (shRNAs) following the protocol of He et al. (*12*). shRNAs used in this study were synthesized *in vitro* (primer sequences provided in Table S1), diluted to 800ng/ul in nuclease-free water (Ambion AM9937) with a final concentration of 0.2 mg/ml Alexa-555 RNAse free dextran (Invitrogen D34679) to facilitate injection. To account for non-specific effects, control embryos were injected with an shRNA (Table S1; 800ng/ul) that was not complimentary to any part of the genome (*42*). Embryos were raised to the early planula stage (72 hours post fertilization) at 16C and effects of knockdowns were assayed via immunofluorescence, *in situ* hybridization, or quantitative PCR (qPCR). Independent confirmation of the effect of ZNF845 knockdown on cnidocyte specification was assayed using a splice-blocking morpholino (GeneTools; Table S1) as previously described (*11*). Briefly, lyophilized morpholinos were reconstituted in nuclease-free water to 1 mM following the manufacturer’s instructions. Before each use, the stock was heated to 60C for 5 mins and centrifuged for ~1 minute before being diluted to a final working concentration of 0.3 mM in nuclease free water with 0.2 mg/ml RNase-free dextran. To control for non-specific effects, ZNF845-injected embryos were compared to embryos injected with a standard control morpholino (GeneTools; Table S1) prepared the same way and injected at the same concentration as the ZNF845 MO. Splicing defects were confirmed using PCR and gel electrophoresis as described previously (*11*).

### Cell and tissue analysis

To assay effects of gene knockdown on cnidocyte specification, developing cnidocytes were labeled, imaged, and counted as described in a similar study (*11*) using an antibody directed against minicollagen4 (Mcol4) (*13*). Additional targets of ZNF845 and NR12 shRNAs were assessed using *in situ* hybridization as previously described (*11*). For qPCR analysis, embryos injected with control shRNA and embryos injected with ZNF845 shRNA were both compared to uninjected embryos raised under the same conditions. Five replicates of each condition (ZNFshRNA, control shRNA, and uninjected/WT) representing five independent injections performed on different days were compared using the delta-delta CT method and the PCR package (43) in the R statistical computing environment (*44*). ZNF845 expression in Figure 1D are presented as fold-change, relative to ZNF845 expression in uninjected embryos and statistical significance for qPCR in Figure 1D was calculated from the comparison of SoxB2 MO and PaxA MO-injected embryos relative to control MO-injected embryos. mRNA expression values in Figures 1F and 2A are presented as fold-change relative to expression of EF1B in the uninjected embryos (arbitrarily set to 1) and significance was calculated from the comparison of ZNF845 shRNA-injected embryos to control shRNA-injected embryos. Cell counts in Figures 1E, 1G, 1I, 2I, 2J, and 2K were analyzed with a Mann-Whitney U nonparametric test for two-way comparisons (control vs target shRNA/MO) and are presented as mean ± standard deviation. Statistical significance for all quantitative comparisons is indicated (*) where p<1E-02.

### Maximum likelihood phylogeny

Phylogenetic analysis of ZF-C_2_H_2_ domains was performed using a modification of a previously published protocol (*45*). In brief, an alignment was generated using a custom script (hmm2aln, available at github.com/josephryan) and the ZF-C_2_H_2_ HMM (PF00096) in predicted proteomes from all target taxa. For cnidarians, we sampled two anthozoans: *Nematostella vectensis* and *Acropora digitifera*, and two medusozoans: *Hydra magnipapillata* and *Nemopilema nomurai* (*46*). For bilaterians, we sampled *Homo sapiens*, *Drosophila melanogaster*, and *Caenorhabditis elegans*. Download information for each taxon is provided in Table S3. The alignment contains over 11,000 ZF-C2H2 domains from these combined taxa and is provided in Data S1. To generate the phylogeny, we first used the model finder function (-MF) with IQTREE to determine the best substitution model (VT+R8) and then generated a single tree and applied 500 bootstraps using fast bootstrapping.

## Acknowledgements and Funding

We are grateful to Namrata Ajuha and Malcolm Moses for their research assistance.

## Funding

This work was funded by the National Aeronautics and Space Administration (grant NNX14AG70G to MQM) and the National Science Foundation (grant 1542597 to JFR).

## Data and Materials Availability

All data is available in the manuscript or the supplementary materials.

## Notes

### Competing Interest Statement

The authors have declared no competing interest.

