## Supplementary Figure S1 for "A novel regulatory gene promotes novel cell fate by suppressing ancestral fate in the sea anemone *Nematostella vectensis*"

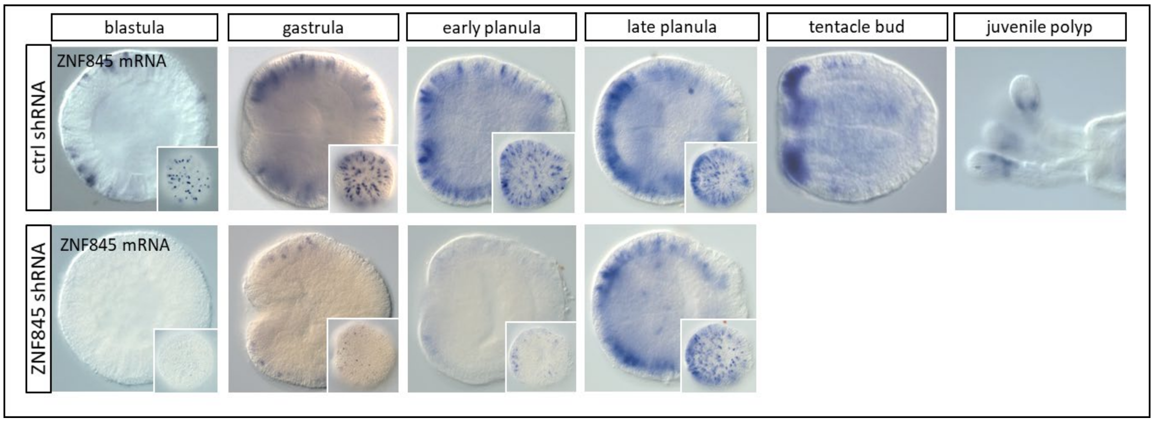


**Fig. S1. Efficacy of ZNF845 shRNA.** ZNF845 mRNA expression in embryos injected with control shRNA (top panels) or ZNF845 shRNA (bottom panels). Nearly complete knockdown of ZNF845 mRNA is shown up to the late planula stage (120hpf at 16C), after which the native expression recovers. Insets show surface detail; the oral pole is oriented to the left in all images.
