## Supplementary Figure S2 for "A novel regulatory gene promotes novel cell fate by suppressing ancestral fate in the sea anemone *Nematostella vectensis*"

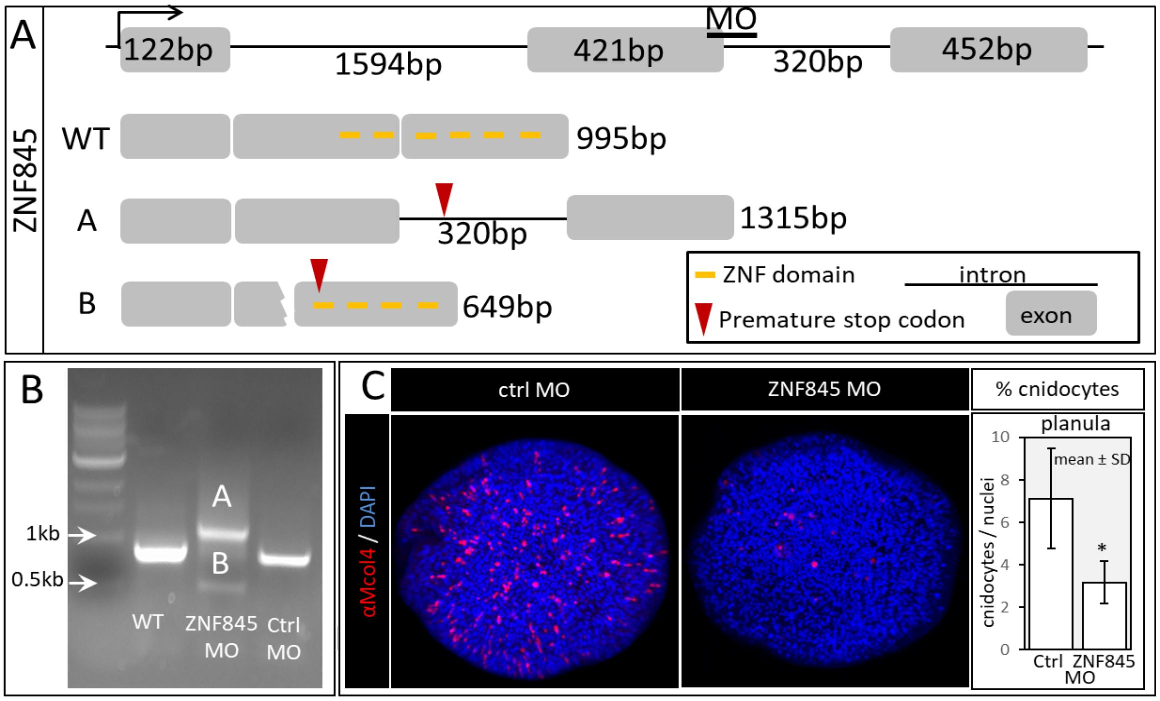


**Fig. S2. Knockdown of ZNF845 with a splice-blocking morpholino phenocopies the results from shRNA knockdowns.** (A) Cartoon representation of the wildtype (WT) and expected splice variants (A and B) resulting from a morpholino (MO) recognizing the exon2/intron2 boundary. (B) Agarose gel showing the WT band for ZNF845 (995bp) for uninjected embryos and embryos injected with standard control (Ctrl) MO. Embryos injected with the ZNF845 MO had two phenotypes: inclusion of the second intron (1315bp band) or mis-splicing of the second and third exons (649bp band). Both splice variants result in premature stop codons eliminating ZNF domains. (C) Immunofluorescent images of early planula stage embryos labeled with DAPI (nuclei, blue) and anti-Mcol4 antibody (cnidocytes, red). (D) Counts of anti-Mcol4 labeled cells relative to the total number of nuclei. Significance (*) is indicated as p<1E-02. See Table S2 for supporting information.
