## Supplementary Figure S3 for "A novel regulatory gene promotes novel cell fate by suppressing ancestral fate in the sea anemone *Nematostella vectensis*"

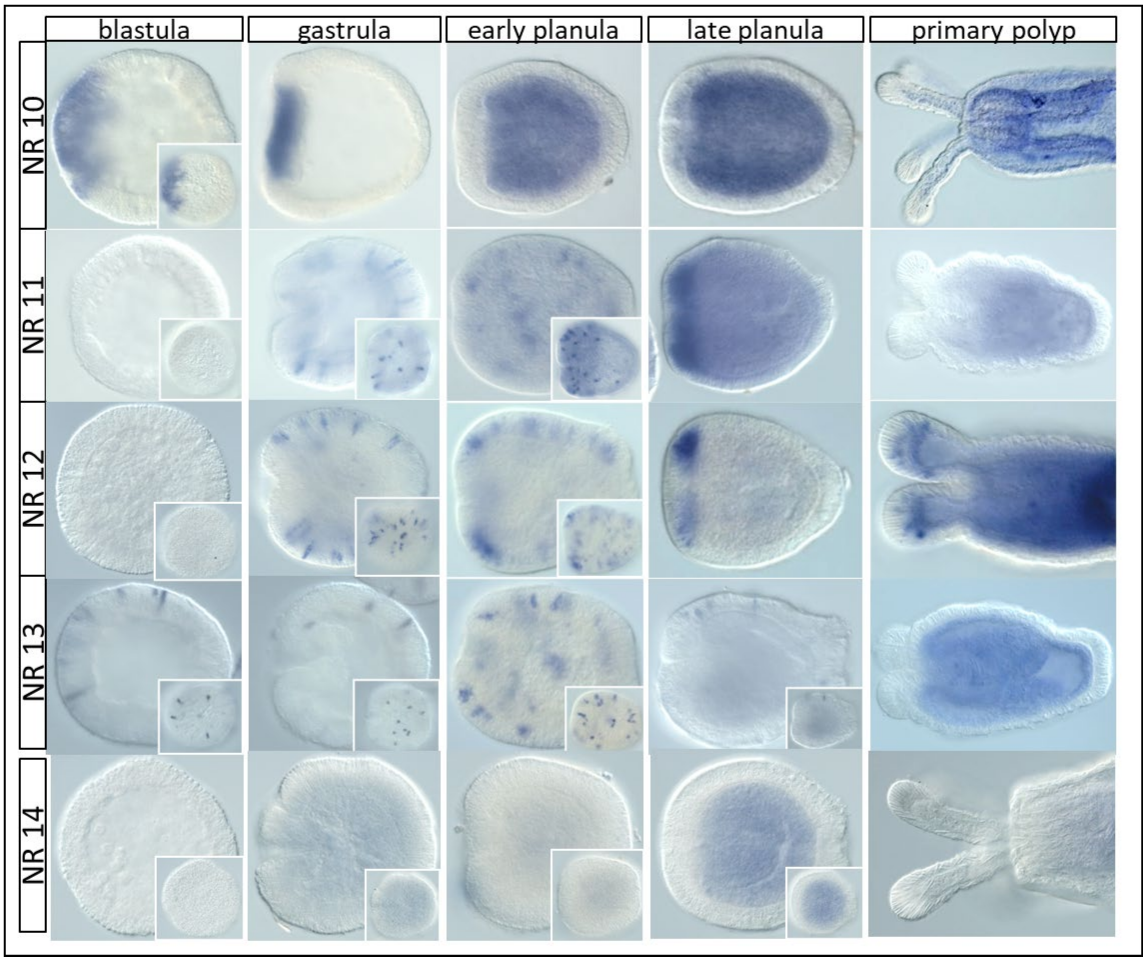


**Fig. S3. Expression of the five NR2F orthologs during development in *N. vectensis*.** NR10 (JGI PID 189134) mRNA is expressed in the presumptive blastopore before the onset of gastrulation and then becomes restricted to the early invaginating endoderm. NR11 (JGI PID 242271), NR12 (JGI PID 165424), and NR13 (JGI PID 203423) are expressed in individual cells throughout the ectoderm. No expression of NR14 (JGI PID 94800) could be detected. The oral pole is oriented to the left in all images; insets show surface detail.
