## Supplementary Figure S4 for "A novel regulatory gene promotes novel cell fate by suppressing ancestral fate in the sea anemone *Nematostella vectensis*"

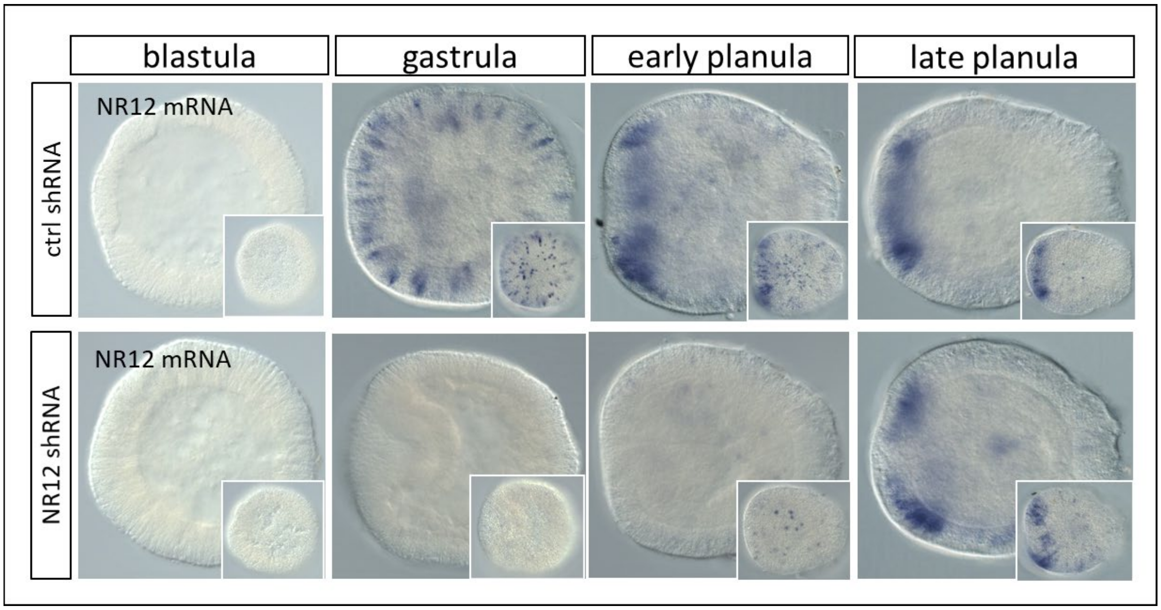


**Fig. S4. Efficacy of NR12 shRNA.** Expression of NR12 mRNA in embryos injected with control shRNA (top panels) or NR12 shRNA (bottom panels). Nearly complete knockdown of NR12 mRNA is seen up to the late planula stage, after which the native expression recovers. The oral pole is oriented to the left in all images; insets show surface detail.
