## Supplementary Figure S5 for "A novel regulatory gene promotes novel cell fate by suppressing ancestral fate in the sea anemone *Nematostella vectensis*"

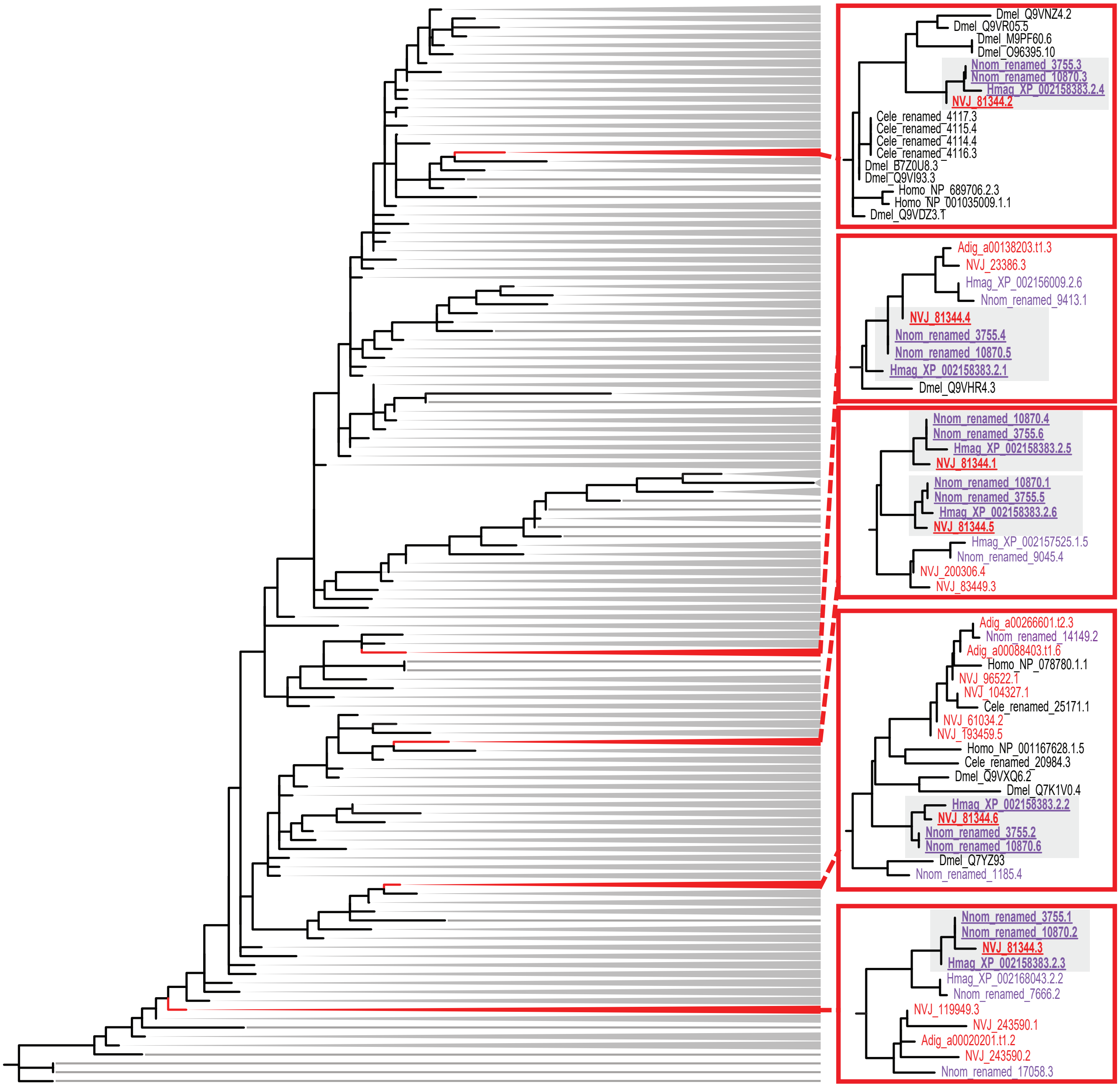


**Fig. S5 Evolutionary history of the ZNF domains in ZNF845.** Maximum likelihood tree of $C_2H_2$ ZNF domains (PF00096) from four cnidarian and three bilaterian taxa. Red boxed insets show regions of detail around each domain from the *N. vectensis* ortholog of ZNF845. Domain IDs from anthozoans are indicated in red, medusozoans are indicated in purple, and bilaterians in black. Domains from the ZNF845 orthologs in each cnidarian lineage are bolded and underlined. Sister relationships were recovered for each of the six ZNF domains from the ZNF845 ortholog in *N. vectensis* (JGI protein ID: 81344), *H. magnipapillata* (NCBI accession: XP_002158383.2), and *N. nomurai* (genome IDs: 3755 and 10870). The bilaterian ZNF domains with the greatest similarity to each of the six ZNF845 domains are found in six different proteins suggesting ZNF845 arose after the divergence of cnidarians and bilaterians from their common ancestor.

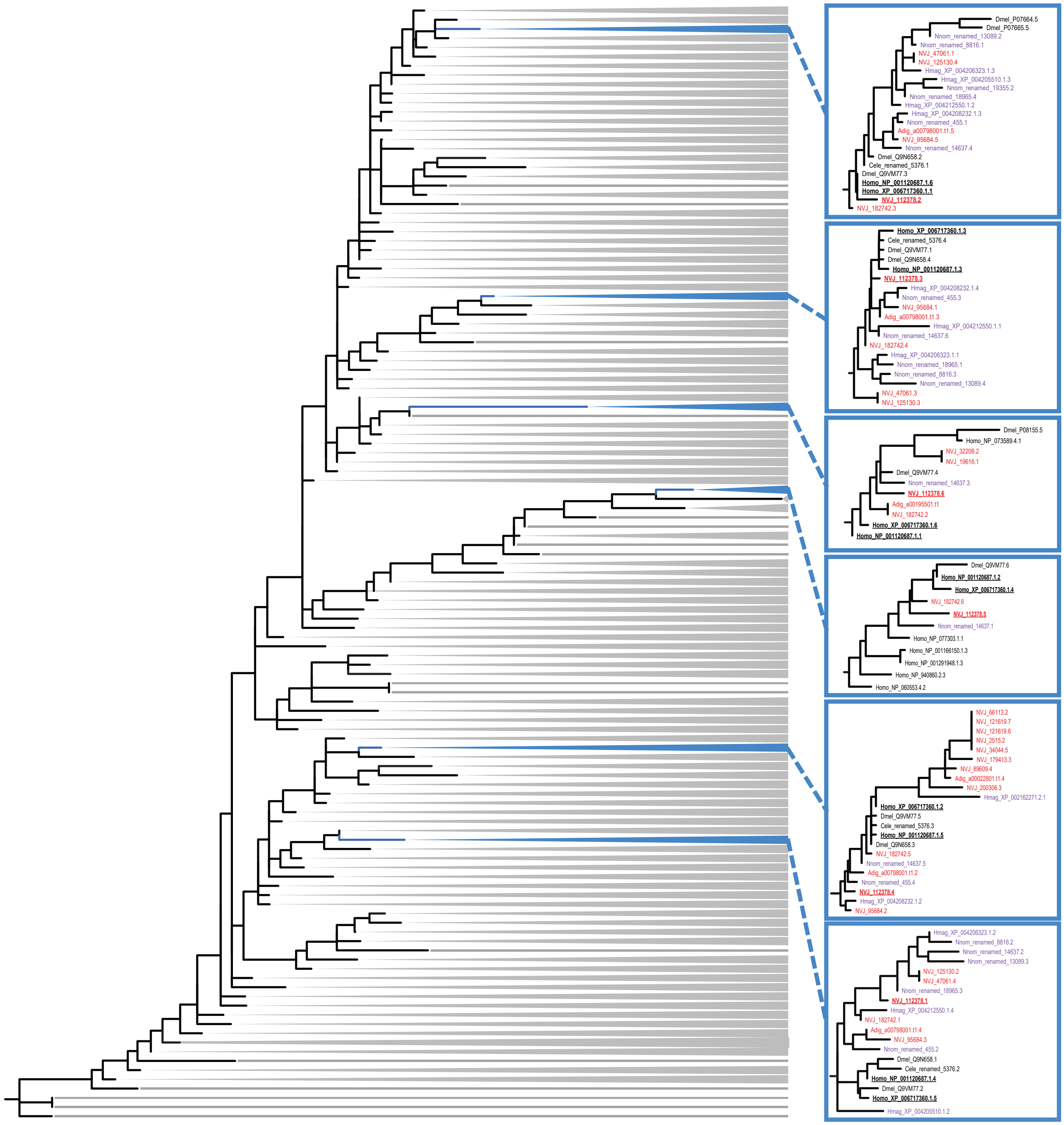


**Fig. S6 Evolutionary history of the $C_2H_2$ ZNF domains in Gfi1B.** The same maximum likelihood tree shown in Fig. S5; blue insets show the details of the regions around each domain from the *N. vectensis* ortholog of Gfi1B. For each domain in this six-domain protein, we recover a sister relationship between the *N. vectensis* ortholog (JGI protein ID: 112378) and the orthologs from *H. sapiens* (NCBI accession: XP_006717360.1, NP_001120687.1) and *D. melanogaster* (Q9VM77). This suggests the six-domain protein was present in the common ancestor of cnidarians and bilaterians.
