## Supplementary Figure S6 for "A novel regulatory gene promotes novel cell fate by suppressing ancestral fate in the sea anemone *Nematostella vectensis*"

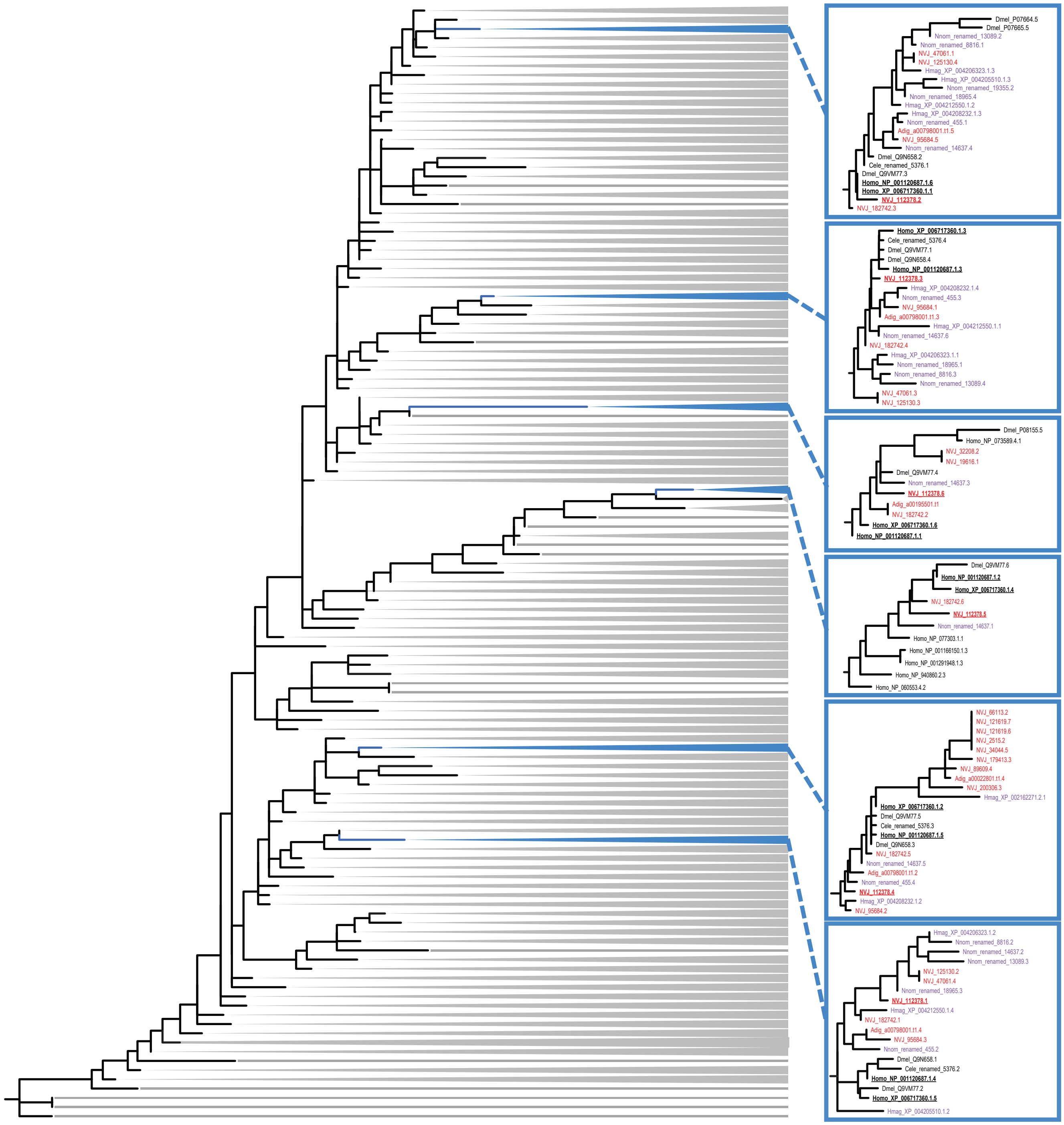


**Fig. S6 Evolutionary history of the $C_2H_2$ ZNF domains in Gfi1B.** The same maximum likelihood tree shown in Fig. S5; blue insets show the details of the regions around each domain from the *N. vectensis* ortholog of Gfi1B. For each domain in this six-domain protein, we recover a sister relationship between the *N. vectensis* ortholog (JGI protein ID: 112378) and the orthologs from *H. sapiens* (NCBI accession: XP_006717360.1, NP_001120687.1) and *D. melanogaster* (Q9VM77). This suggests the six-domain protein was present in the common ancestor of cnidarians and bilaterians.
